# Early Brain Imaging can Predict Autism: Application of Machine Learning to a Clinical Imaging Archive

**DOI:** 10.1101/471169

**Authors:** Gajendra J. Katuwal, Stefi A. Baum, Andrew M. Michael

## Abstract

A comprehensive investigation of early brain alterations in Autism Spectrum Disorder (ASD) is critical for understanding the neuroanatomical underpinnings of autism and its early diagnosis. Most previous brain imaging studies in ASD, however, are based on children older than 6 years – well after the average age of ASD diagnosis (~46 months). In this study, we use brain magnetic resonance images that were collected as part of clinical routine from patients who were later diagnosed with ASD. Using 15 ASD subjects of age three to four years and 18 age-matched non-ASD subjects as controls, we perform comprehensive comparison of different brain morphometric features and ASD vs. non-ASD classification by Random Forest machine learning method. We find that, although total intracranial volume (TIV) of ASD was 5.5 % larger than in non-ASD, brain volumes of many other brain areas (as a percentage of TIV) were smaller in ASD and can be partly attributed to larger (>10 %) ventricles in ASD. The larger TIV in ASD was correlated to larger surface area and increased amount of cortical folding but not to cortical thickness. The white matter regions in ASD had less image intensity (predominantly in the frontal and temporal regions) suggesting myelination deficit. We achieved 95 % area under the ROC curve (AUC) for ASD vs. non-ASD classification using all brain features. When classification was performed separately for each feature type, image intensity yielded the highest predictive power (95 % AUC), followed by cortical folding index (69 %), cortical and subcortical volume (69 %), and surface area (68 %). The most important feature for classification was white matter intensity surrounding the rostral middle frontal gyrus and was lower in ASD (d = 0.77, p = 0.04). The high degree of classification success indicates that the application of machine learning methods on brain features holds promise for earlier identification of ASD. To our knowledge this is the first study to leverage a clinical imaging archive to investigate early brain markers in ASD.

## 1 Introduction

According to a recent CDC survey (CDC, 2014), 1 in 68 children have a diagnosis of Autism Spectrum Disorder (ASD) and a more recent CDC survey of parents indicated that this number can be as high as 1 in 45 (Zablotsky et al., 2015) Early detection of ASD is important as it allows for the application of early intervention methods. Early intervention is effective in reducing the impact of impairments (Dawson et al., 2010) and may result in more positive long-term outcomes for the child (Rogers and Vismara, 2010; Pickles et al., 2016). The risk for autism is influenced by genetic and pre, peri, and post-natal environmental factors (Sandin et al., 2014). The environmental factors related to the risk for autism can interact with genetic factors making the identification of ASD etiology extremely difficult. In addition to better clinical outcomes, early detection of ASD can also help in separating the effects of post-natal environmental risk factors of ASD and this can lead to improving our knowledge of its etiology.

The median age of ASD diagnosis is 46 months (CDC, 2014) and current ASD diagnosis is based on a clinical assessment of the individual’s behavior and intellectual abilities. However, this approach is limited as early diagnosis is not straightforward and behaviorally based diagnosis procedures can be subjective, time consuming, and inconclusive due to factors such as comorbidity (Close et al., 2012). Furthermore, such an approach does not provide insight into the neural underpinnings of ASD since it is based only on behavioral symptoms. Magnetic resonance imaging (MRI) is a non-invasive tool widely used to capture brain morphology. ASD diagnosis based on MRI can be objective and can potentially be utilized even at the prenatal and neonatal stage (Glenn, 2010) and hence can be a useful tool for brain biomarker discovery for early detection of ASD. MRI has already been successfully utilized for early diagnosis of brain disorders such as Alzheimer’s (Frisoni et al., 2010; Hampel et al., 2010; Hazlett et al., 2017).

A number of MRI studies have reported alterations in brain regions involved in language and social behavior in ASD, particularly in fronto-temporal regions (Bigler et al., 2007; Ha et al., 2015), and the amygdala-hippocampus complex (Groen et al., 2010; Nordahl et al., 2012). Early brain overgrowth (Campbell et al., 2014) and alterations in corpus callosum (Wolff et al., 2015) and cerebellum (D’Mello et al., 2015) have also been reported. However, these findings have been somewhat inconsistent (Jumah et al., 2016; Katuwal et al., 2016b). Similarly, in recent years, there have been a number of studies that have applied machine learning techniques for ASD vs. non-ASD classification using MRI derived brain features. These studies have reported high classification accuracies (>80%) for well-matched datasets (Ecker et al., 2010; Jiao et al., 2010; Katuwal et al., 2015). Most MRI studies that investigate ASD brain alterations have used subjects older than six years and there are only a very few studies in early childhood (< four years) (Nordahl et al., 2012; Auzias et al., 2014; Padilla et al., 2015; Hazlett et al., 2017). Of the studies using subjects less than age four years, only a small subset of brain features such as volume, total cortical surface area, and shape have been investigated. As such, there is a significant knowledge gap in brain imaging research on early ASD brain alterations. A comprehensive investigation of brain alterations in early childhood in children with ASD is needed to better characterize this disorder and merits special attention due to the importance of early detection of ASD.

In this study, we compare brain features of ASD subjects in early childhood (three to four years) with non-ASD subjects of the same age group using a comprehensive set of cortical and sub-cortical morphometric features. In addition, we include image intensity features (mean and standard deviation of intensity of brain structures). Multi-variate brain alterations are identified in a purely data-driven approach by applying machine learning models trained for ASD vs. non-ASD classification.

## 2 Material and Methods

### 2.1 Subjects

Head MR images used in this study were obtained from the clinical imaging archive of Geisinger Health System, Danville, PA. We queried the Geisinger imaging archive to identify ASD patients who had an MR image before the age of 4 and identified 167 patients. We then found 437 age matched subjects who did not have ASD. The flowchart of subject selection is presented as Figure A2 in Appendix A. We performed a strict data quality analysis to remove images with imaging artifacts, motion, lesions, and abnormally large ventricles. The remaining images (247 non-ASD and 122 ASD) were processed using Freesurfer v 5.3.0 (Fischl, 2012) recon-all workflow with default settings and the images that failed Freesurfer segmentation (5 non-ASD and 7 ASD) were excluded from the study. We then removed 120 non-ASD subjects as they had a neurodevelopmental disorder as identified by ICD 9 codes. This resulted a total of 112 non-ASD and 115 ASD subjects. To avoid gender confounds (Lai et al., 2013), 54 female subjects (20 non-ASD, 34 ASD) were excluded.

**Table 1:** Subject Demographics & Image Parameters

|  | **ASD** | **Control** | **D** | **p-value** |
| --- | --- | --- | --- | --- |
| **N** | 15 | 18 |  |  |
| **Age (months)** | 42.6 ± 3.5 (37.7 to 47.3) | 41.4 ± 2.9 (37.0 to 47.0) | 0.23 | 0.76 |
| **Echo time (ms)** | 5.7 ± 1.1 | 5.0 ± 1.7 | 0.24 | 0.81 |
| **Flip angle** | 23.7 ± 8.8 | 24.6 ± 8.9 | 0.16 | 0.99 |
| **Slice spacing** | 1.4 ± 0.4 | 1.5 ± 0.4 | 0.19 | 0.96 |
| **Slice thickness** | 1.8 ± 0.2 | 1.8 ± 0.3 | 0.15 | 1 |
| **Voxel size x (mm)** | 0.6 ± 0.2 | 0.6 ± 0.2 | 0.25 | 0.75 |
| **Voxel size y(mm)** | 0.6 ± 0.2 | 0.6 ± 0.2 | 0.25 | 0.75 |
| **Scanner Model** | 9 Gx 5 Gxt 1 Pv | 5 Gx 3 Gxt 3 Gxc 3 Pv 4 NA | 0.25 | 0.75 |

**D:** Two-sample Kolmogorov-Smirnov statistic **Gx:** GE Medical Systems Signa hdx **Gxt:** GE Medical Systems signa hdx **Gxc:** Medical Systems Signa Excite **Pv:** Philips Medical Systems Achieva **NA:** Not Available

The lower threshold for age was based on the applicability of Freesurfer on brain images of very young subjects. Freesurfer which was initially designed for adult subjects may not provide reliable estimates for subjects younger than three years. As images of subjects as young as three years have been successfully segmented by Freesurfer (Retico et al., 2016), lower threshold for age was set to three years. The upper threshold for age was set to four years to look for early markers.

The segmentation results from Freesurfer of remaining 41 images (23 non-ASD, 18 ASD) were assessed using the ENIGMA protocols (http://enigma.ini.usc.edu/protocols/imaging-protocols/). Five non-ASD and one ASD images with imperfect Freesurfer segmentations were excluded from the study. Our final dataset consisted of 15 ASD subjects(42.6 ± 3.5 months; 37.7 to 47.3 months) and 18 non-ASD (41.4 ± 2.9 months; 37.0 to 47.0 months) subjects which was used as the control group; no significant age difference (Kolgmogorov-Smirnov statistic = 0.23, p = 0.76). See Table 2.1 for subject demographics and image parameters.

### 2.2 Brain Features Extraction

Brain features were extracted using the recon-all workflow of Freesurfer v. 5.3.0 (Dale et al., 1999; Fischl et al., 1999, 2002; Ségonne et al., 2004). The volume of 40 sub-cortical structures from Aseg atlas (Fischl et al., 2002) were extracted. Similarly, volume, surface area, Gaussian curvature, mean curvature, folding index, curvature index, thickness mean, thickness standard deviation (std.), intensity mean, and intensity std. of 34 cortical structures from Desikan-Killiany atlas (Desikan et al., 2006) were extracted. In total, 687 brain morphometric and intensity features were derived for each image. Intensity mean and intensity std. of a brain structure is the mean and standard deviation respectively of the voxel intensities within the brain structure. The volume features of each subject were normalized by its TIV since relative volumes can be directly compared across subjects and are more robust against scanner effects (Takao et al., 2011). To eliminate the effects of image intensity biases, intensity features of each image were normalized by their respective sums across all the brain structures.

### 2.3 Univariate Analysis

The statistical significance of ASD vs. non-ASD brain feature differences was calculated using two sample t-tests. For each feature type, multiple comparisons correction was performed using false discovery rate (Benjamini and Hochberg, 1995). The effect size (ES) of the differences were quantified by Cohen’s d (Baron-Cohen et al., 2000).

### 2.4 Multivariate Analysis: ASD vs. non-ASD classification using Random Forest

ASD vs. non-ASD classification was performed by Random Forest (RF) (Breiman, 2001) classification models trained with all 687 FreeSurfer brain features. At first RF models were trained using all brain features and then models for each feature type were trained separately. Classification success was quantified by area under the ROC curve (AUC). The average classification success across the data was estimated by 5-fold cross-validation with stratified folds. Similarly, classification contribution or importance scores for features were average across the 5 folds. For detailed methodology of multivariate analysis, please see Appendix A at the end.

## 3 Results

### 3.1 ASD vs. non-ASD Brain Feature Differences

The brain features for which ASD vs. non-ASD differences were statistically significant (p < 0.05, uncorrected) are presented in Figure 1. Effect sizes of all these differences were moderate to large (Cohen’s d > 0.5). None of the features survived the multiple comparisons correction and all the results reported heron are before the correction.

**Figure 1:**
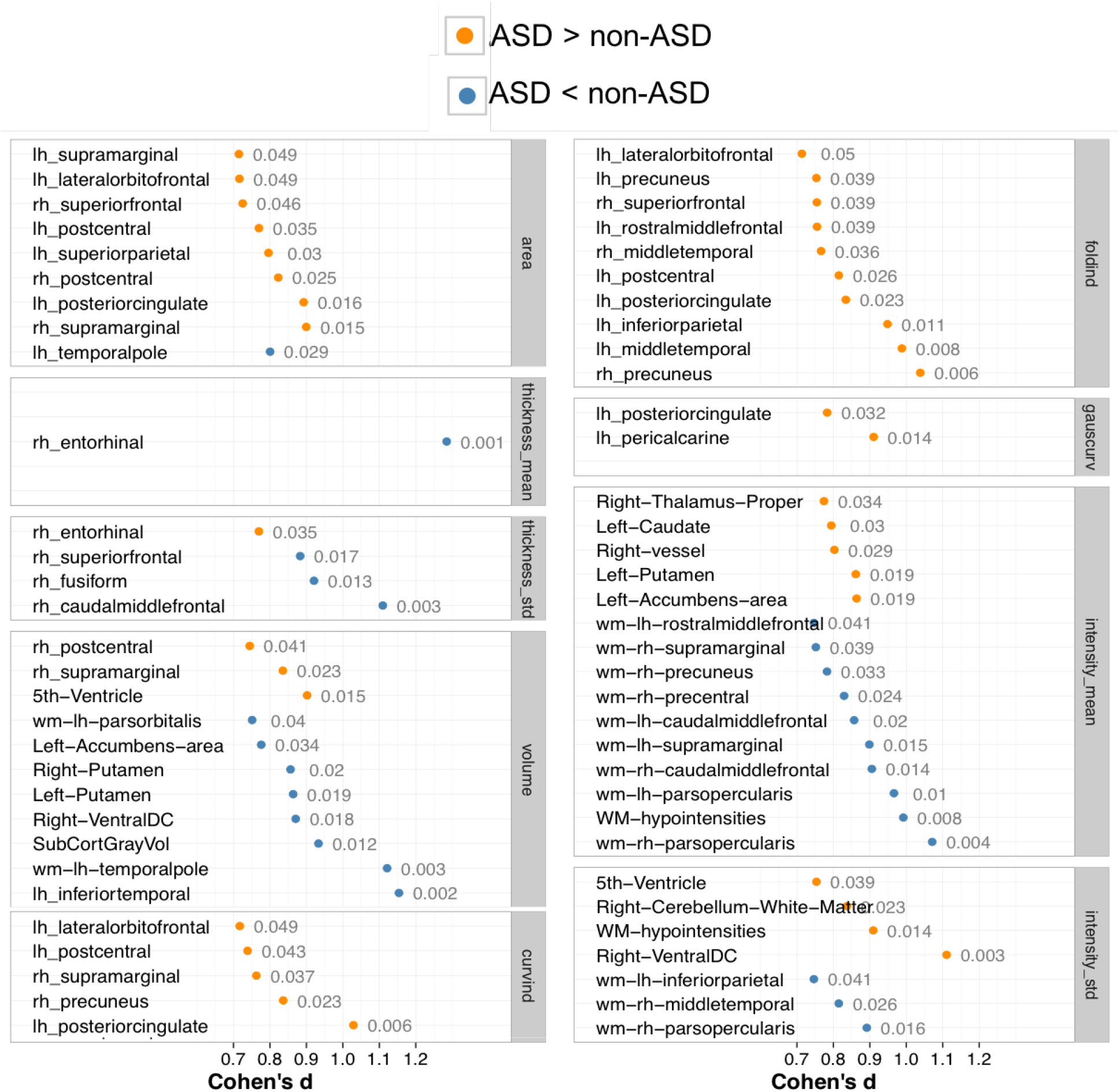
Effect sizes (Cohen’s d) of brain features with statistically significant ASD vs. non-ASD differences (at p =0.05, uncorrected for multiple comparison). Effect sizes are denoted by orange dots for ASD > non-ASD and by blue dots for non-ASD > ASD. The numeric text near the dots denote p-values. Volume features were normalized by total intracranial volume.

Statistically significant area features were mainly located in frontal, temporal, supramarginal, and posterior cingulate regions. Except for the area of the left temporal pole, the other eight statistically significant area features were larger in ASD. The thickness mean of entorhinal gyrus was larger in ASD and the effect size was large (d > 1; p = 0.0009) whereas the thickness std. of the right entorhinal gyrus was smaller in ASD (d = 0.75; p = 0.035). Similarly, thickness std. of the right superior frontal, fusiform, and caudal middle frontal gyri were larger in ASD (d > 0.60).

Most of the volumes with statistically significant differences (not corrected for multiple comparisons) were smaller in ASD except for the right postcentral, supramarginal, and 5*_th_* ventricle volumes. However, it should be noted these reported volume features have been normalized by TIV. Raw volumes of these structures were in fact larger in ASD but the differences were not statistically significant. Similarly, all global raw volumes were larger in ASD, but when normalized by TIV, most of them were smaller in ASD. This discrepancy is mainly because TIV in ASD was larger than in non-ASD by 5.5% (d = 0.4, p = 0.17); see Figure B1 in Appendix B. Larger TIV in ASD was mainly due to the larger ventricles. All ventricles were larger (>10%) in ASD, in particular the 5th ventricle was 290% larger in ASD (p = 0.015). Total ventricular CSF volume was 27.89% larger in ASD (d = 0.38, p = 0.28) and after normalizing by TIV, it was 19.14% larger (d = 0.29, p = 0.42).

The folding index and curvature index features with statistically significant differences were present in the frontal, temporal, cingulate, postcentral, and precuneus regions and all were larger in ASD. Folding indices of 58 out of 68 cortices were higher in ASD. The differences in curvature index of the left posterior cingulate gyrus and the folding index of the right precuneus gyrus were large (d > 1). Similarly, the Gaussian curvature of the left posterior cingulate gyrus and left pericalcarine gyri were larger in ASD.

In image intensity mean features, there were two distinct type of differences: in general, white matter (WM) regions had lower and sub-cortical regions had higher mean intensity in ASD. Mean intensity of the WM structures near frontal, supramarginal, precuneus, precentral, and pars opercularis gyri were lower in ASD. In contrast, mean intensity of the right thalamus, left caudate, left putamen, and left accumbens were higher in ASD. In general, in ASD, intensity std. were higher in WM near gyri but smaller in cerebellum WM and corpus callosum (not significant).

### 3.2 ASD vs. non-ASD Classification

ASD vs. non-ASD classification AUC scores for each feature type are presented in Figure 2. The point and error bar represent the mean and standard deviation of the AUC scores from 5 test folds. An average AUC of 0.92 was achieved when all brain features were used. When the classification was performed separately for each brain feature type, we found that most of the predictive power came from the intensity mean (AUC = 0.83), folding index (AUC=0.69), volume (AUC=0.69), and area (AUC = 0.69) features. When intensity mean and intensity std. features were used together, AUC of 0.95 was achieved.

**Figure 2:**
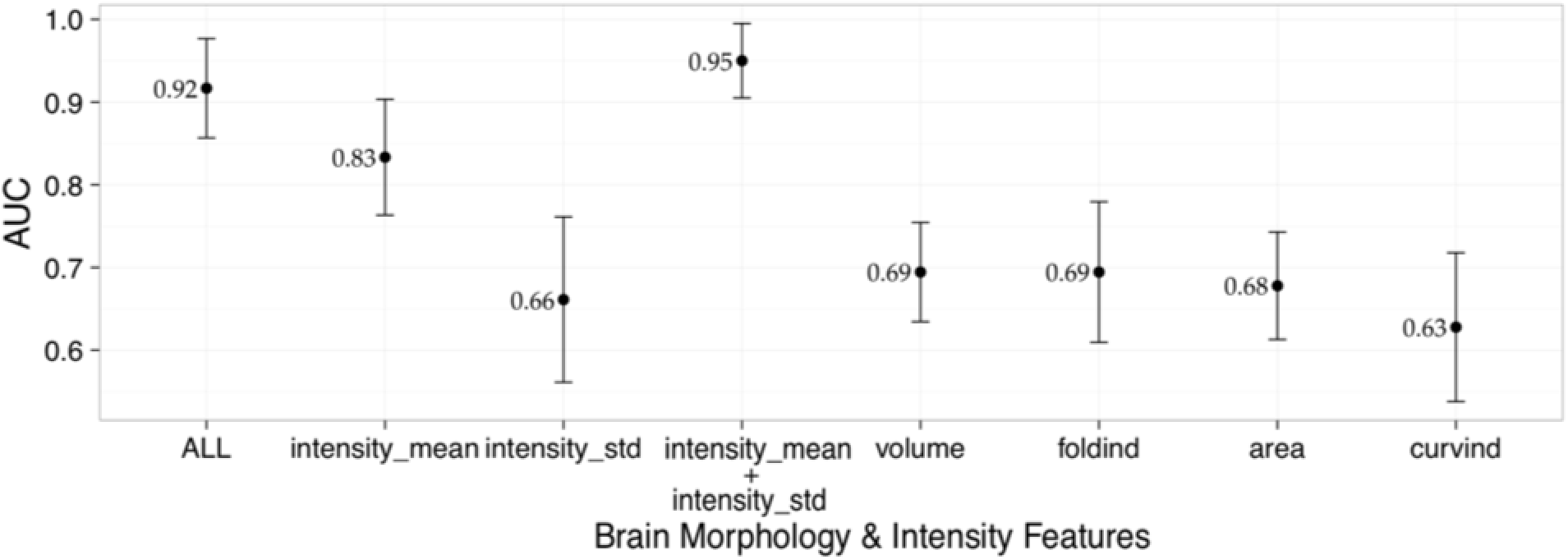
ASD vs. non-ASD classification AUC scores. Mean classification AUC for different types of brain features across five folds of testing. The error bar represents the standard deviation of the AUC scores of 5 folds.

### 3.3 Important Features for Classification

The important features in classification for the feature types that yielded high AUCs (intensity mean, folding index, volume, and area) are presented in Figure 3. After sorting the importance scores of the features in descending order, the features whose cumulative sum of scores was at least 50% of the total importance scores were deemed as important for classification and are presented in Figure 3. In Figure 3, a feature is represented by a bar and its length is proportional to the contribution of the feature towards classification relative to the most important feature. The numbers at the left of the bars represent the effect size of the ASD vs. non-ASD difference and the asterisk represent statistical significance of the difference at 0.05.

**Figure 3:**
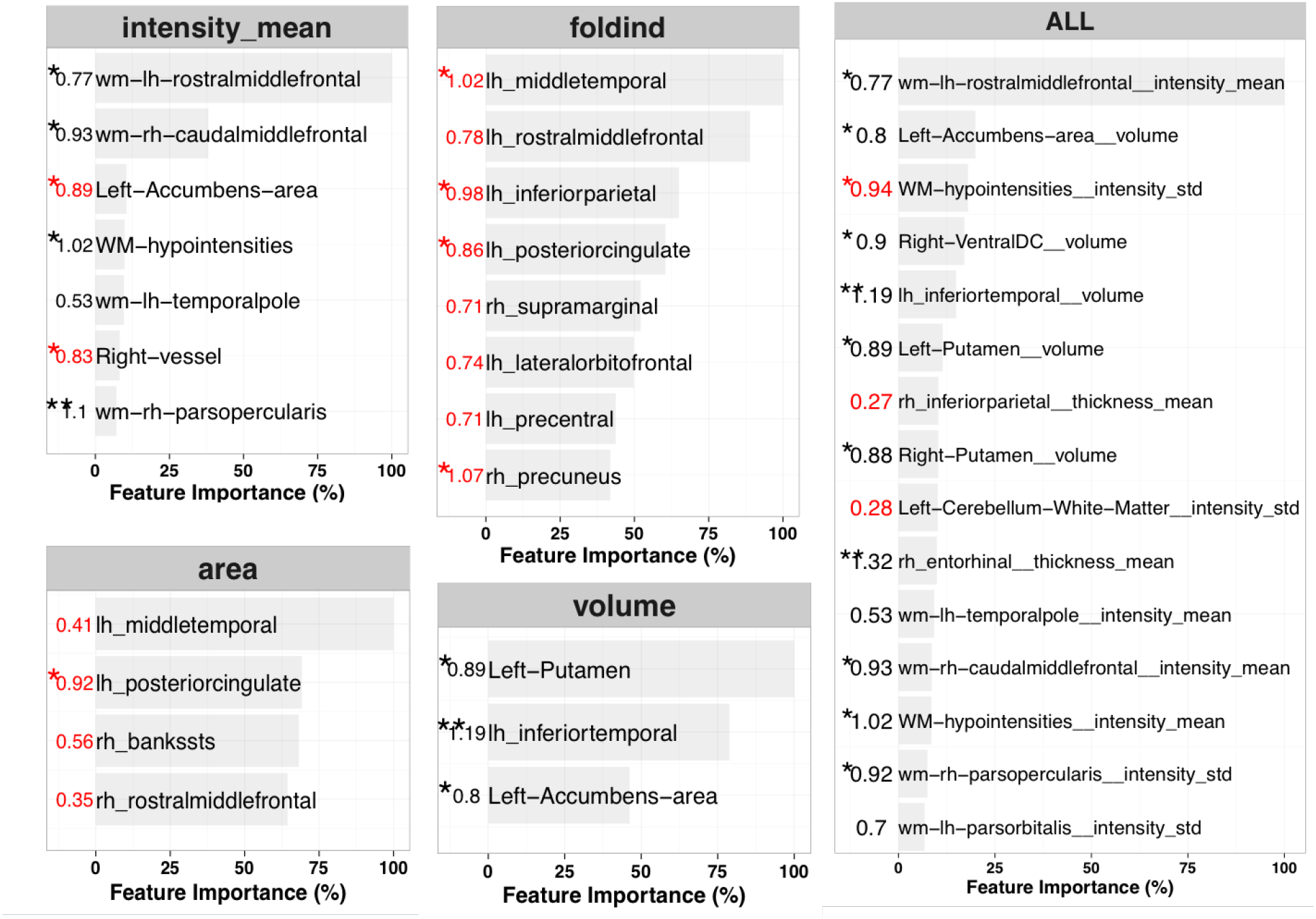
Important features for ASD vs. non-ASD classification. The length of a horizontal bar represents feature importance as a percentage of the most important feature in the same feature category. The numbers at the left of the bars represent ASD – non-ASD Cohen’s d (positive is represented by red) and the asterisk represents statistical significance at 0.05 from two sample t-test.

When only image intensity mean features were used for classification, the most important feature was the intensity mean of the WM neighboring rostral middle frontal gyrus and compared to non-ASD, it was lower in ASD (d = 0.77, p = 0.04). The mean intensities of the WM neighboring the right caudal middle frontal, left temporal pole, and the right pars opercularis were also important for classification and were smaller in ASD. Other important brain structures were the left accumbens-area, WM-hypointensities, and right vessel. Similarly, when only area features were used for classification, the surface areas of the left middle temporal gyrus, left posterior cingulate gyrus, right bank of superior temporal sulcus (bankssts), and right rostral middle frontal gyrus were important for classification. All of the important area features were larger in ASD. In classification using only folding index features, the left middle temporal gyrus was the most important brain structure for classification. It was also the most important structure while performing classification using area features. Other important structures were left middle temporal, left rostral middle frontal, and left inferior parietal gyrus. All of the important folding index features were larger in ASD. Similarly, when classification was performed only with volume features, the volumes of the left putamen, left inferior temporal gyrus, and left accumbens-area were important for classification and these features were smaller in ASD after normalizing for TIV. When all the features were used for classification, the most important feature was the mean intensity of the WM neighboring the rostral middle frontal gyrus; note that the same region was the most important feature in the classification using image intensity features only. The mean intensities of WM connecting left temporal pole and right caudal middle frontal gyrus, were important for classification and were smaller in ASD.

## 4 Discussion

We achieved high ASD vs. non-ASD classification success using brain morphometric and image intensity features. The important features for classification were mainly present in the frontal and temporal regions and these regions have been previously associated with ASD (Bigler et al., 2007; Ha et al., 2015). Most of the discriminative or predictive power for classification came from the intensity features followed by folding index, volume, and area. In summary, compared to non-ASD three main brain alterations in ASD were noted: (1) abnormally larger ventricles, (2) higher cortical gyrification, and (3) lesser WM intensity in frontal and temporal region. This is the first study to perform ASD vs. non-ASD classification in a very young population (3 to 4 years) using comprehensive brain features and to achieve high classification success rates. Very few studies have performed similar investigations to identify brain alterations in ASD.

### 4.1 Early brain overgrowth in ASD

One of the most replicated findings in ASD is that toddlers with ASD (age 2-4 years) on average have a larger head size than non-ASD (Carper et al., 2002; Courchesne et al., 2011; Hazlett et al., 2012; Campbell et al., 2014). A recent study by (Hazlett et al., 2017) also reported 5.5% larger TIV in high risk ASD compared to negative high risk ASD at the age of two years. Our study also shows 5.5% larger TIV in ASD. The larger TIV in ASD in our study was mainly due to the larger ventricular volume in ASD (27.9% and 19.1% as a percentage of TIV) whereas volume of other brain regions (as a percentage of TIV) were smaller in ASD. Padilla et al. (2015) have also reported smaller brain volumes in temporal, occipital, insular, and limbic regions in ASD after adjusting for total brain volume.

### 4.2 Overgrowth is related to the increase in surface area and is relatively independent to cortical thickness

In our study, only one thickness feature (right entorhinal gyrus thickness, smaller in ASD) had statistically significant group difference. However, there were nine area features (eight were larger in ASD) whose differences were statistically significant and cortical surface area was 7% larger in ASD. In addition, thickness features yielded near chance classification success suggesting low discriminative power for ASD classification while area features yielded AUC of 0.69. Two major longitudinal studies (Hazlett et al., 2011, 2017) using subjects of age similar to our study report similar findings. A study by Hazlett et al. (2011) found no differences in cortical thickness but larger surface area in ASD at both 2 years and 4.5 years. Similarly, a recent study by Hazlett et al. (2017) using subjects of age 6 to 24 months, report 7% larger cortical surface area in ASD but no difference in cortical thickness. They also report insignificant contribution of thickness features in classification of ASD subjects with ASD from non-ASD subjects. These results suggests that while the group difference in cortical surface area increases with the early brain overgrowth, the difference in cortical thickness remains somewhat constant.

### 4.3 Relationship between overgrowth, cortical expansion, and higher cortical folding in ASD brains

Folding indices of most of the gyri (58 out of 68) were greater in ASD and folding index features yielded AUC of 0.69. On average, the cortex of ASD brains were 12.7% more folded than non-ASD. This suggests that ASD brains generally exhibit more gyrification and cortical folding. Similar to our study, Ecker et al. (2016) have reported that ASD individuals had a significant increase in gyrification around the left pre and post-central gyrus. Similarly, a study by Auzias et al. (2014) reported higher folding of the right intraparietal, the left medial frontal, and the left central sulci in children with ASD.

#### 4.3.1 Hyper-expansion of the cortex during early brain overgrowth is a cause of higher cortical folding in ASD

The higher amount of cortical folding in ASD may be the after effect of early brain overgrowth in ASD toddlers. Based on insights from the cortical gyrification hypothesis based on cortical expansion (Tallinen et al., 2014, 2016), we hypothesize that higher folding in ASD could be due to larger compressive stress in the cortex produced by the tangential hyper-expansion of the cortical layer. We found that the correlation between the average amount of cortical folding and total surface area of the cortex 0.68 (p = 0.005) in ASD and 0.78 (p = 0.0002) in non-ASD. Similarly, the correlation between the average amount of cortical folding and TIV was 0.53 (p = 0.04) in ASD and 0.65 (p = 0.03) in non-ASD.

#### 4.3.2 Larger ventricles contribute to higher cortical folding in ASD by creating additional compressive stress

We also hypothesize that the abnormally larger ventricles in ASD induces additional compressive stress in the cortex and this leads to higher cortical folding. In this study, we note that the correlation between average amount of cortical folding and total ventricular CSF was 0.56 (p = 0.02) for ASD, but was −0.13 (p = 0.6) for non-ASD. The statistically significant positive correlation in ASD suggests the possibility of some degree of cortical folding may be due to the larger ventricles. In contrast, the non-significant correlation in TDC suggests that the cortical folding is not affected by the ventricles because they are of normal size and hence do not extend additional compressive stress to the cortex.

#### 4.3.3 Larger volume of extra-axial fluid in ASD contributes to higher cortical folding in ASD

Furthermore, it is possible that the larger volume of extra-axial fluid (CSF in the sub-arachnoid space) in ASD also contributes to the compressive stress in the cortex and hence more folding. A study by Shen et al. (2013) has reported that extra-axial fluid in ASD was 25% more than low-risk typical infants at the age of 6 to 24 months. In our study, we could not perform the analysis for the extra-axial fluid volume because FreeSurfer does not output the measure since T1 MRI does not have sufficient contrast between CSF and air as both appear dark in T1.

### 4.4 WM in frontal and temporal regions less myelinated in ASD

Image intensity mean of the WM neighboring frontal and temporal regions were lower in ASD and some of these features were the most important for classification. Less image intensity mean in WM means it is less bright suggesting less myelination surrounding the axons. Models using brain intensity features yielded high classification success and most of the important features were WM neighboring frontal and temporal regions. This suggests myelination deficits in frontal and temporal regions may be a potential early brain marker of ASD.

Several previous studies have also reported WM myelination deficits in ASD (Peters et al., 2012; Zinkstok et al., 2012; Croteau-Chonka et al., 2016). Zinkstok et al. (2012) reported that individuals with ASD had significantly less myelin content in numerous brain regions and WM tracts. In addition, they report that the observed myelination deficit increased with autism severity. Similarly, Peters et al. (2012) have reported myelination deficits in WM in autistic brains of age 0.5 to 25 years. These results suggest that brain disconnectivity resulting from insufficient development of the myelin sheath may be one of the underlying causes of ASD and myelination deficits in frontal and temporal regions in particular, may be a potential marker for early detection of ASD.

### 4.5 Limitations of this study

#### 4.5.1 Small sample size

The sample size of the study was only 33. Further studies with larger sample sizes are needed to evaluate the robustness of our findings.

#### 4.5.2 Findings are based on only male subjects

Only male subjects were used in this study to exclude gender confounds. Findings of this study can be interpreted in the context of brain abnormalities in male ASD subjects only and not the general ASD population because previous studies have shown gender differences in ASD brain morphometry.

#### 4.5.3 Effect of preprocessing methods

Freesurfer was used to extract morphological features from brain MRIs. There have been several studies showing the effects of preprocessing methods and in some cases the bias introduced by the methods were larger than the actual group differences (Katuwal et al., 2016a). One way to minimize the possibility of interpreting method effects as biological effects is to use multiple preprocessing methods and to cross-validate the results. However, at the time of this study, Freesurfer was the only available tool that generated the extensive morphological brain features we wanted to use for machine learning. In addition, Freesurfer which was initially designed for adult subjects may introduce some biases during the preprocessing of the images of the young subjects who are three to four years old. However, images of subjects as young as three years have been successfully segmented by Freesurfer (Retico et al., 2016) and this is why we decided to use the images of the subjects as young as three years.

### 4.6 Lack of first-hand confirmation of ASD/non-ASD status

ASD status and non-ASD status were based on diagnostic codes from the electronic health records. Although, strict research criteria were not applied to confirm ASD status, each subject had at least 4 occurrences of ASD diagnoses assigned by clinical experts.

### 4.7 Effect of IQ

We have no information on IQ for either group. Significant group mean differences in IQ or differences in IQ distribution could be responsible for some of the findings (rather than ASD status).

### 4.8 Non-ASD subjects were not typically developing children

The non-ASD subjects which were used as controls were imaged for some reason and may have had atypical development. As such, results presented here are brain differences between ASD and non-ASD subjects and not ASD and typically developing children.

## 5 Conclusion

We were able to classify ASD from non-ASD subjects with high accuracy, using MRI derived brain features. We identified five potential brain markers for early detection of ASD: larger ventricles, larger total brain volume, larger cortical surface area, higher cortical folding, and myelination deficits particularly in frontal and temporal regions. Further, we were able to show that higher cortical folding in ASD brains may be an after effect of early brain overgrowth and the additional compression in the cortex due the abnormally large ventricles.

## Appendix A: Materials and Methods

Random forest (RF) (Breiman, 2001) is an ensemble of decision tree classifiers and its output class is the popular class decided by the voting of its member classifiers. Area under the ROC curve (AUC) was used to measure the success of classification and the average classification success was estimated by 5-fold cross-validation with stratified folds. Scikit-learn 0.17.1 (Pedregosa and Varoquaux, 2011) was used to perform all the multivariate analyses. For more information on Random Forest, see. Information gain measure was used to measure the quality of a node split while growing decision trees. Following hyper parameters were used for optimal model selection — max_depth, the number of features used to judge the quality of the impurity of a node, min_samples_split, the minimum number of samples required in a node for further splitting, and min_samples_leaf, the minimum number of samples required to be at a leaf node. Hyper parameter optimization was performed through random search.

**Figure A1:**
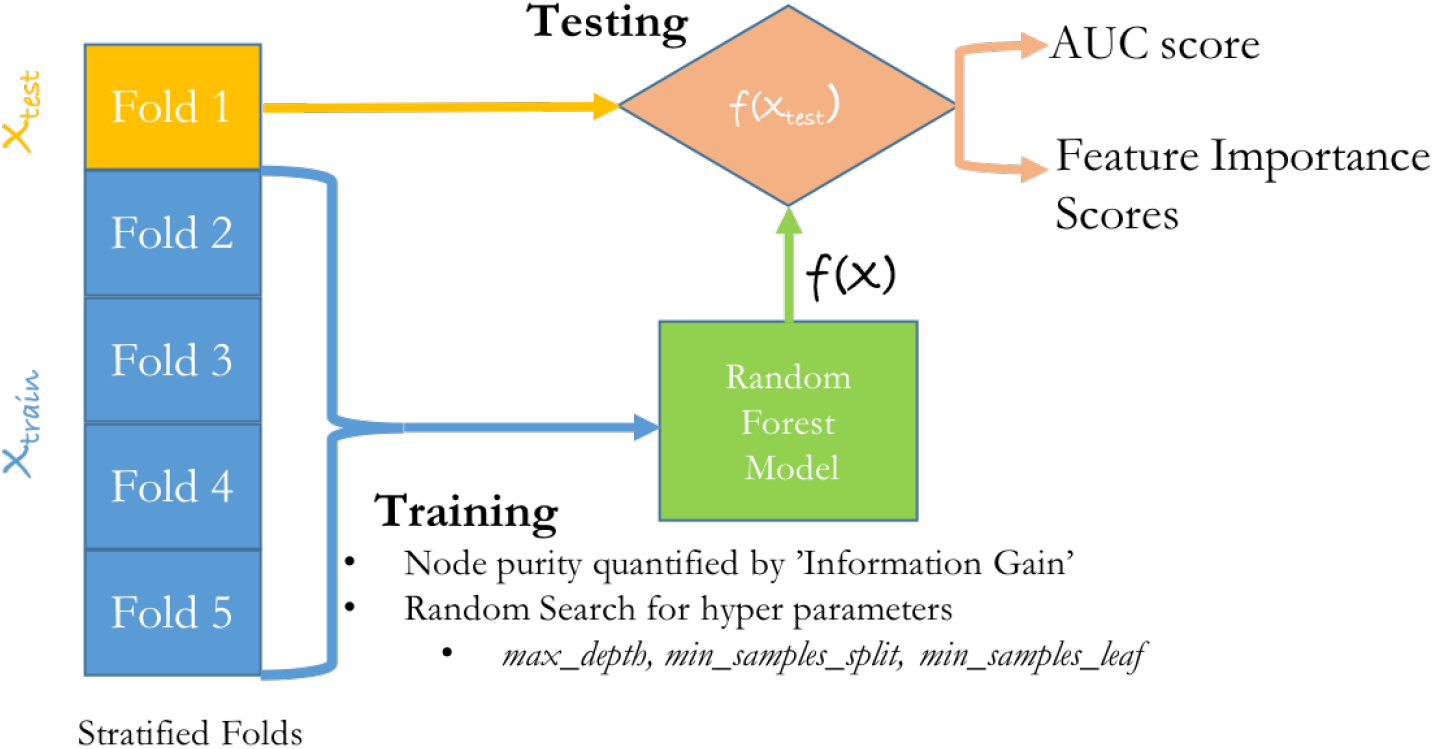
Classification flowchart. Independent experiments were performed for each feature type. For each classification experiment, the model success was quantified by AUC metric and the average model assessment was estimated by 5–fold cross-validation. The 5-folds were generated by stratified sampling to ensure dissimilarity between the folds. Each classification was performed using random forest classifier where the node purity was quantified by ‘Information Gain’. The optimum hyper parameters (max_depth, min_samples_split, min_samples_leaf) were estimated using random search. Feature importance scores across the 5 folds were averaged.

**Figure A2:**
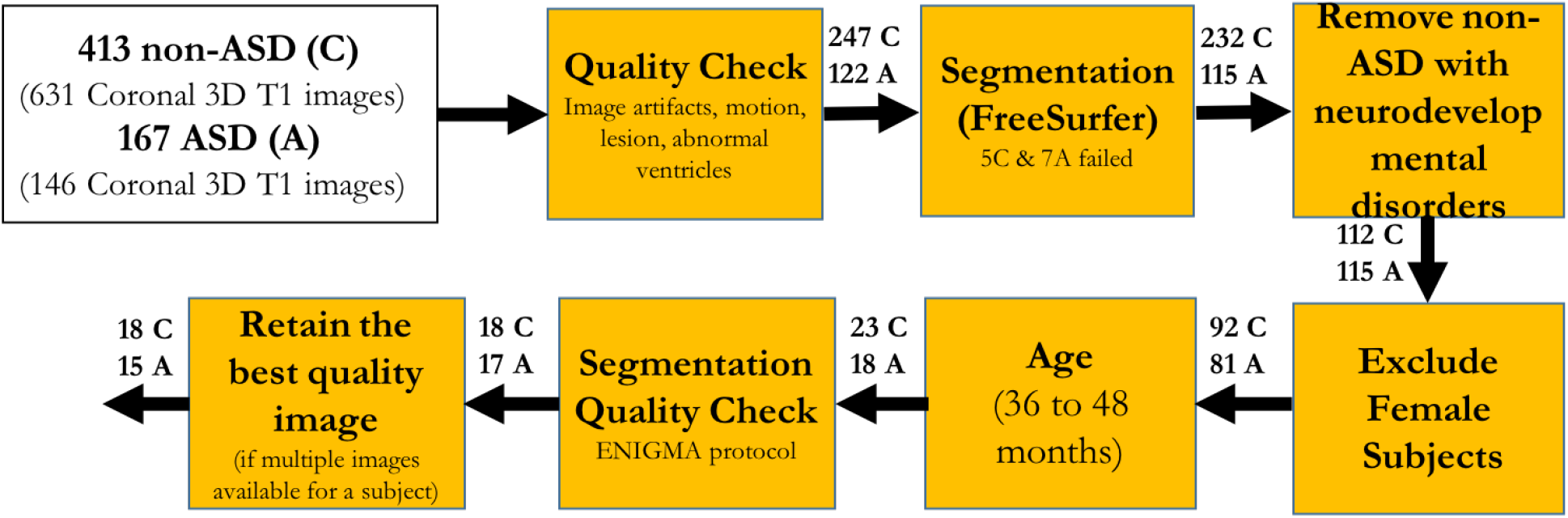
Subject Selection Flowchart.

## Appendix B: Supplementary Results

**Figure B1:**
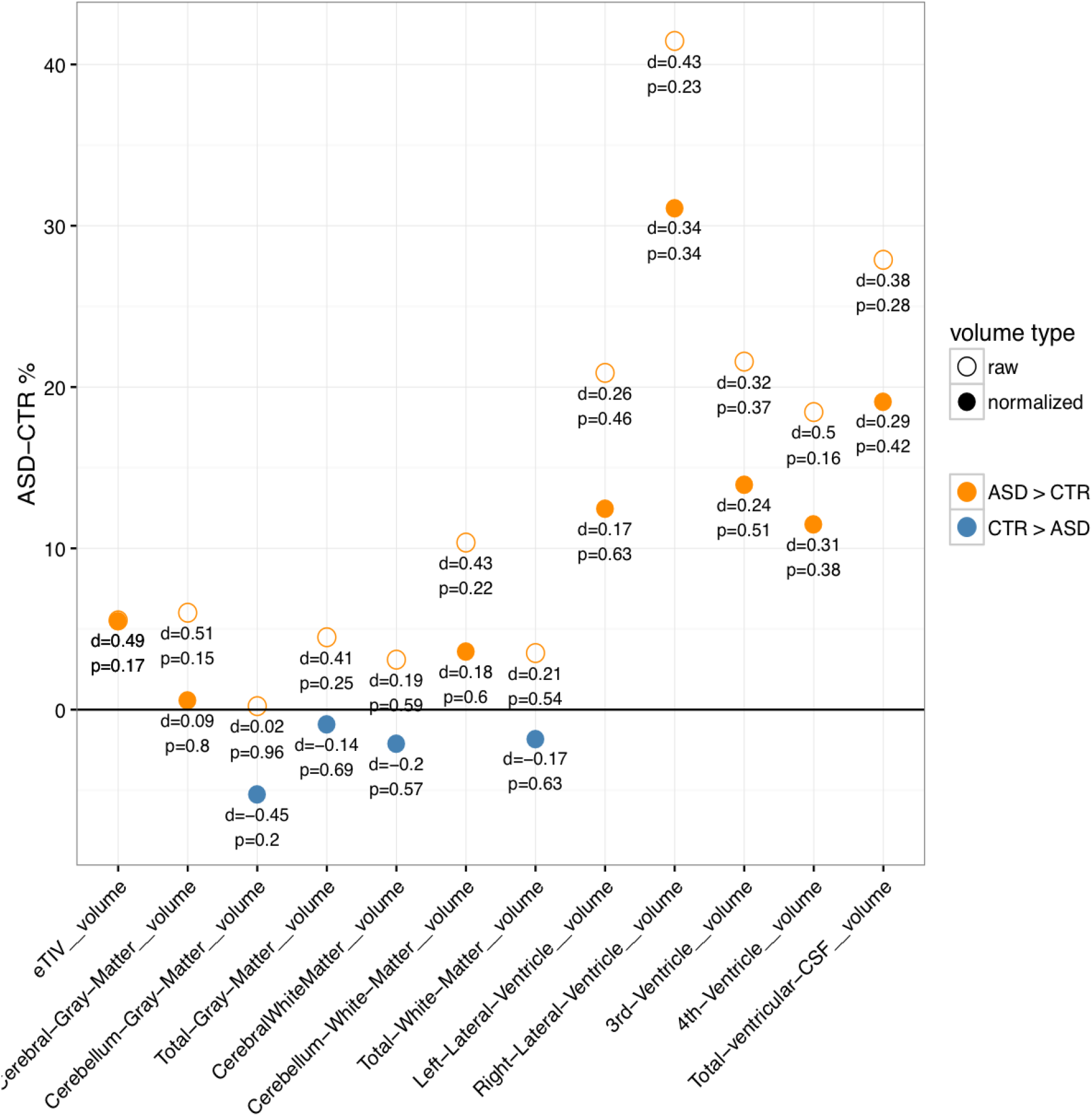
ASD versus non-ASD (CTR) global brain volume differences. Total inter-cranial volume (TIV) was larger in ASD and was mainly due to abnormally large ventricles. Normalized global volumes were smaller in ASD.

## Footnotes

Maybe we should share all the feature differences as an excel file?

Generally refered as septum pallidum http://www.radlex.org/RID/RID6525.

http://scikit-leam.org/stable/modules/generated/sklearn.ensemble.RandomForestClassifler.html

## Acknowledgement

The authors would like to thank Chao Zhang, Chase Doughtery, and Viraj Adduru for their help in quality check of the brain images used in this study.

## References

Auzias, G., Viellard, M., Takerkart, S., Villeneuve, N., Poinso, F., Fonséca, D. D., Girard, N., and Deruelle, C. (2014). Atypical sulcal anatomy in young children with autism spectrum disorder. NeuroImage: Clinical, 4:593–603.

Baron-Cohen, S., Ring, H. a., Bullmore, E. T., Wheelwright, S., Ashwin, C., and Williams, S. C. R. (2000). The amygdala theory of autism. Neuroscience and Biobehavioral Reviews, 24(3):355–364.

Benjamini, Y. and Hochberg, Y. (1995). Controlling the false discovery rate: a practical and powerful approach to multiple testing.

Bigler, E. D., Mortensen, S., Neeley, E. S., Ozonoff, S., Krasny, L., Johnson, M., Lu, J., Provencal, S. L., McMahon, W., and Lainhart, J. E. (2007). Superior temporal gyrus, language function, and autism. Developmental neuropsychology, 31(2):217–38.

Breiman, L. (2001). Random Forrest. Machine Learning, pages 1–33.

Campbell, D. J., Chang, J., and Chawarska, K. (2014). Early generalized overgrowth in autism spectrum disorder: Prevalence rates, gender effects, and clinical outcomes. Journal of the American Academy of Child and Adolescent Psychiatry, 53(10):1063–1073.

Carper, R. A., Moses, P., Tigue, Z. D., and Courchesne, E. (2002). Cerebral lobes in autism: early hyperplasia and abnormal age effects. NeuroImage, 16(4):1038–51.

CDC (2014). Prevalence of autism spectrum disorder among children aged 8 years – autism and developmental disabilities monitoring network, 11 sites, United States, 2010. Technical Report 2.

Close, H. a., Lee, L.-C., Kaufmann, C. N., and Zimmerman, a. W. (2012). Co-occurring Conditions and Change in Diagnosis in Autism Spectrum Disorders. Pediatrics, 129(2):e305–e316.

Courchesne, E., Karns, C. M., Davis, H. R., Ziccardi, R., Carper, R. a., Tigue, Z. D., Chisum, H. J., Moses, P., Pierce, K., Lord, C., Lincoln, A. J., Pizzo, S., Schreibman, L., Haas, R. H., Akshoomoff, N. a., and Courchesne, R. Y. (2011). Unusual brain growth patterns in early life in patients with autistic disorder: an MRI study. Neurology, 57(24):2111–2111.

Croteau-Chonka, E. C., Dean, D. C., Remer, J., Dirks, H., O’Muircheartaigh, J., and Deoni, S. C. L. (2016). Examining the relationships between cortical maturation and white matter myelination throughout early childhood. NeuroImage, 125:413–421.

Dale, A. M., Fischl, B., and Sereno, M. I. (1999). Cortical surface-based analysis. I. Segmentation and surface reconstruction. NeuroImage, 9(2):179–94.

Dawson, G., Rogers, S., Munson, J., Smith, M., Winter, J., Greenson, J., Donaldson, A., and Varley, J. (2010). Randomized, controlled trial of an intervention for toddlers with autism: The early start denver model. Pediatrics, 125(1):e17–e23.

Desikan, R. S., Ségonne, F., Fischl, B., Quinn, B. T., Dickerson, B. C., Blacker, D., Buckner, R. L., Dale, A. M., Maguire, R. P., Hyman, B. T., Albert, M. S., and Killiany, R. J. (2006). An automated labeling system for subdividing the human cerebral cortex on MRI scans into gyral based regions of interest. NeuroImage, 31(3):968–80.

D’Mello, A. M., Crocetti, D., Mostofsky, S. H., and Stoodley, C. J. (2015). Cerebellar gray matter and lobular volumes correlate with core autism symptoms. NeuroImage: Clinical, 7:631–639.

Ecker, C., Andrews, D., Dell’Acqua, F., Daly, E., Murphy, C., Catani, M., Thiebaut De Schotten, M., Baron-Cohen, S., Lai, M. C., Lombardo, M. V., Bullmore, E. T., Suckling, J., Williams, S., Jones, D. K., Chiocchetti, A., and Murphy, D. G. M. (2016). Relationship between cortical gyrification, white matter connectivity, and autism spectrum disorder. Cerebral Cortex, 26(7):3297–3309.

Ecker, C., Marquand, A., Mourão-Miranda, J., Johnston, P., Daly, E. M., Brammer, M. J., Maltezos, S., Murphy, C. M., Robertson, D., Williams, S. C., and Murphy, D. G. M. (2010). Describing the brain in autism in five dimensions–magnetic resonance imaging-assisted diagnosis of autism spectrum disorder using a multiparameter classification approach. The Journal of neuroscience : the official journal of the Society for Neuroscience, 30(32):10612–10623.

Fischl, B. (2012). FreeSurfer. NeuroImage, 62(2):774–81.

Fischl, B., Salat, D. H., Busa, E., Albert, M., Dieterich, M., Haselgrove, C., van der Kouwe, A., Killiany, R., Kennedy, D., Klaveness, S., Montillo, A., Makris, N., Rosen, B., and Dale, A. M. (2002). Whole brain segmentation: automated labeling of neuroanatomical structures in the human brain. Neuron, 33(3):341–55.

Fischl, B., Sereno, M. I., Tootell, R. B., and Dale, a. M. (1999). High-resolution intersubject averaging and a coordinate system for the cortical surface. Human brain mapping, 8(4):272–84.

Frisoni, G. B., Fox, N. C., Jack, C. R., Scheltens, P., and Thompson, P. M. (2010). The clinical use of structural MRI in Alzheimer disease. Nature Reviews, 6(2):67–77.

Glenn, O. A. (2010). MR imaging of the fetal brain. Pediatric Radiology, 40(1):68–81.

Groen, W., Teluij, M., Buitelaar, J., and Tendolkar, I. (2010). Amygdala and Hippocampus Enlargement During Adolescence in Autism. Journal of the American Academy of Child & Adolescent Psychiatry, 49(6):552–560.

Ha, S., Sohn, I.-J., Kim, N., Sim, H. J., and Cheon, K.-A. (2015). Characteristics of Brains in Autism Spectrum Disorder: Structure, Function and Connectivity across the Lifespan. Experimental Neurobiology, 24(4):273–284.

Hampel, H., Frank, R., Broich, K., Teipel, S. J., Katz, R. G., Hardy, J., Herholz, K., Bokde, A. L. W., Jessen, F., Hoessler, Y. C., Sanhai, W. R., Zetterberg, H., Woodcock, J., and Blennow, K. (2010). Biomarkers for Alzheimer’s disease: academic, industry and regulatory perspectives. Nature reviews. Drug discovery, 9(7):560–574.

Hazlett, H. C., Gu, H., Munsell, B. C., Kim, S. H., Styner, M., Wolff, J. J., Elison, J. T., Swanson, M. R., Zhu, H., Botteron, K. N., Collins, D. L., Constantino, J. N., Dager, S. R., Estes, A. M., Evans, A. C., Fonov, V. S., Gerig, G., Kostopoulos, P., Mckinstry, R. C., Pandey, J., Paterson, S., Pruett Jr, J. R., Schultz, R. T., Shaw, D. W., Zwaigenbaum, L., and Piven, J. (2017). Early brain development in infants at high risk for autism spectrum disorder. Nature Publishing Group, 542(7641):348–351.

Hazlett, H. C., Poe, M., Gerig, G., Styner, M., Chappell, C., Smith, R. G., Vachet, C., and Piven, J. (2011). Early Brain Overgrowth in Autism Associated with an Increase in Cortical Surface Area Before Age 2 years. Archives of General Psychiatry, 68(5):467–476.

Hazlett, H. C., Poe, M. D., Lightbody, A. A., Styner, M., MacFall, J. R., Reiss, A. L., and Piven, J. (2012). Trajectories of early brain volume development in fragile X syndrome and autism. Journal of the American Academy of Child and Adolescent Psychiatry, 51(9):921–933.

Jiao, Y., Chen, R., Ke, X., Chu, K., Lu, Z., and Herskovits, E. H. (2010). Predictive models of autism spectrum disorder based on brain regional cortical thickness. NeuroImage, 50(2):589–599.

Jumah, F., Ghannam, M., Jaber, M., Adeeb, N., and Tubbs, R. S. (2016). Neuroanatomical variation in autism spectrum disorder: A comprehensive review. Clinical Anatomy, 29(4):454–465.

Katuwal, G. J. (2017). Machine Learning Based Autism Detection Using Brain Imaging. PhD thesis, Rochester Institute of Technology.

Katuwal, G. J., Baum, S. A., Cahill, N. D., Dougherty, C. C., Evans, E., Evans, D. W., Moore, G. J., and Michael, A. M. (2016a). Inter-method discrepancies in brain volume estimation may drive inconsistent findings in autism. Frontiers in Neuroscience, 10(SEP):439.

Katuwal, G. J., Baum, S. A., Cahill, N. D., and Michael, A. M. (2016b). Divide and Conquer: Sub-Grouping of ASD Improves ASD Detection Based on Brain Morphometry. PloS one, 11(4):e0153331.

Katuwal, G. J., Cahill, N. D., Baum, S. A., and Michael, A. M. (2015). The Predictive Power of Structural MRI in Autism Diagnosis. In Engineering in Medicine and Biology Society (EMBC), 2015 37th Annual International Conference of the IEEE, pages 4270–4273.

Lai, M.-C., Lombardo, M. V., Suckling, J., Ruigrok, A. N. V., Chakrabarti, B., Ecker, C., Deoni, S. C. L., Craig, M. C., Murphy, D. G. M., Bullmore, E. T., and Baron-Cohen, S. (2013). Biological sex affects the neurobiology of autism. Brain : a journal of neurology, 136(Pt 9):2799–815.

Nordahl, C. W., Scholz, R., Yang, X., Buonocore, M. H., Simon, T., Rogers, S., and Amaral, D. G. (2012). Increased Rate of Amygdala Growth in Children Aged 2 to 4 Years With Autism Spectrum Disorders. 69(1):53–61.

Padilla, N., Eklöf, E., Mårtensson, G. E., Bölte, S., Lagercrantz, H., and Ådén, U. (2015). Poor Brain Growth in Extremely Preterm Neonates Long Before the Onset of Autism Spectrum Disorder Symptoms. Cerebral Cortex, (February 2016):1–8.

Peters, J. M., Sahin, M., Vogel-Farley, V. K., Jeste, S. S., Nelson, C. A., Gregas, M. C., Prabhu, S. P., Scherrer, B., and Warfield, S. K. (2012). Loss of White Matter Microstructural Integrity Is Associated with Adverse Neurological Outcome in Tuberous Sclerosis Complex. Academic Radiology, 19(1):17–25.

Pickles, A., Le Couteur, A., Leadbitter, K., Salomone, E., Cole-Fletcher, R., Tobin, H., Gammer, I., Lowry, J., Vamvakas, G., Byford, S., Aldred, C., Slonims, V., McConachie, H., Howlin, P., Parr, J. R., Charman, T., and Green, J. (2016). Parent-mediated social communication therapy for young children with autism (PACT): long-term follow-up of a randomised controlled trial. The Lancet, 388(10059):2501–2509.

Retico, A., Gori, I., Giuliano, A., Muratori, F., and Calderoni, S. (2016). One-class support vector machines identify the language and default mode regions as common patterns of structural alterations in young children with autism spectrum disorders. Frontiers in Neuroscience, 10(JUN).

Rogers, S. J. and Vismara, L. a. (2010). Evidence-Based Comprehensive Treatments for Early Autism. Journal of Clinical Child and Adolescent Psychology, 37(1):8–38.

Sandin, S., Lichtenstein, P., Larsson, H., Cm, H., and Reichenberg, A. (2014). The familial risk of autism. 311(17):24794370.

Ségonne, F., Dale, a. M., Busa, E., Glessner, M., Salat, D., Hahn, H. K., and Fischl, B. (2004). A hybrid approach to the skull stripping problem in MRI. NeuroImage, 22(3):1060–75.

Shen, M. D., Nordahl, C. W., Young, G. S., Wootton-Gorges, S. L., Lee, A., Liston, S. E., Harrington, K. R., Ozonoff, S., and Amaral, D. G. (2013). Early brain enlargement and elevated extra-axial fluid in infants who develop autism spectrum disorder. Brain, 136(9):2825–2835.

Takao, H., Hayashi, N., and Ohtomo, K. (2011). Effect of scanner in longitudinal studies of brain volume changes. Journal of magnetic resonance imaging : JMRI, 34(2):438–44.

Tallinen, T., Chung, J. Y., Biggins, J. S., and Mahadevan, L. (2014). Gyrification from constrained cortical expansion. Proceedings of the National Academy of Sciences of the United States of America, 111(35):12667–72.

Tallinen, T., Chung, J. Y., Rousseau, F., Girard, N., Lefèvre, J., and Mahadevan, L. (2016). On the growth and form of cortical convolutions. Nature Physics, 476(7358):57–62.

Wolff, J. J., Gerig, G., Lewis, J. D., Soda, T., Styner, M. A., Vachet, C., Botteron, K. N., Elison, J. T., Dager, S. R., Estes, A. M., Hazlett, H. C., Schultz, R. T., Zwaigenbaum, L., and Piven, J. (2015). Altered corpus callosum morphology associated with autism over the first 2 years of life. Brain, 138(7):2046–2058.

Zablotsky, B., Black, L. I., Maenner, M. J., Schieve, L. A., and Blumberg, S. J. (2015). Estimated Prevalence of Autism and Other Developmental Disabilities Following Questionnaire Changes in the 2014 National Health Interview Survey. National Health Statistics Reports, (87):1–21.

Zinkstok, J., Kolind, S., D’Almeida, V., Shahidiani, A., Williams, S. C., Murphy, D. G., and Deoni, S. C. (2012). Is Myelin Content Altered In Young Adults with Autism? In INSAR, Toronto.

